# The microbiome wants what it wants: microbial evolution overtakes experimental host-mediated indirect selection

**DOI:** 10.1101/706960

**Authors:** Jigyasa Arora, Margaret Mars Brisbin, Alexander S. Mikheyev

## Abstract

Microbes ubiquitously inhabit animals and plants, often affecting their host’s phenotype. As a result, even in a constant genetic background, the host’s phenotype may evolve through indirect selection on the microbiome. ‘Microbiome engineering’ offers a promising novel approach for attaining desired host traits but has been attempted only a few times. Building on the known role of the microbiome on development in fruit flies, we attempted to evolve earlier eclosing flies by selecting on microbes in the growth media. We carried out parallel evolution experiments in no- and high-sugar diets by transferring media associated with fast-developing fly lines over the course of four rounds of selection. In each round, we used sterile eggs from the same inbred population, and assayed fly mean eclosion times. Ultimately, flies eclosed seven to twelve hours earlier, depending on the diet, but selection had no effect. 16S sequencing showed that the microbiome did evolve, particularly in the no sugar diet, with an increase in alpha diversity over time. Thus, while microbiome evolution did affect host eclosion times, these effects were incidental. Instead, any experimentally enforced selection effects were swamped by independent microbial evolution. These results imply that selection on host phenotypes must be strong enough to overcome other selection pressures simultaneously operating on the microbiome. The independent evolutionary trajectories of the host and the microbiome may limit the extent to which indirect selection on the microbiome can ultimately affect host phenotype. Random-selection lines accounting for independent microbial evolution are essential for experimental microbiome engineering studies.

## Introduction

Communities of microbes living on or in multicellular host organisms interact with their hosts in diverse ways that often influence host phenotype and fitness (Zilber-rosenberg & Rosenberg 2008). Such host-microbe interactions have traditionally been investigated by experimentally comparing hosts raised without a microbiome (axenic) to hosts inoculated with known components of the microbiome (gnotobiotic) or that receive microbiome transplants composed of complex communities (Turnbaugh et al. 2009). Observations of atypical phenotypes in axenic organisms indicate hosts are dependent on their microbiome and cannot function normally without it. The integral role of the microbiome in shaping host phenotype suggests that desirable host traits can be indirectly selected through microbiome engineering (Mueller & Sachs 2015; Gopal & Gupta 2016; Oyserman et al. 2018). To achieve this, microbes or microbial communities correlated with desired host traits are selected, but selection success is evaluated by measuring host traits (Mueller et al. 2016). This novel approach has numerous practical applications, such as better probiotic design and improved crop yields, in genetically homogeneous or otherwise unaltered hosts.

While applying or administering specific bacterial strains or communities (i.e. probiotics) to achieve a desired host effect is now widespread, true microbiome engineering studies remain rare. Diverse examples of successful probiotic studies include: increased biomass and antioxidant capacity in plants inoculated with *Agrobacterium* (Chihaoui et al. 2015), reduced white pox disease in corals that received a probiotic cocktail of 13 bacterial strains isolated from coral mucus (Alagely et al. 2011), and intestinal epithelial cells with increased ability to keep pathogens from escaping the intestinal tract in mice that were administered *Lactobacillus* strains (Mack et al. 2003). While often successful, probiotic approaches typically rely on relatively simple manipulations of the microbiome by introducing known and culturable bacterial species. Additionally, probiotic studies usually take advantage of some prior knowledge of host-microbe interactions involving the host or microbe(s) of interest. However, microbiomes as communities are more complex than what is generally applied experimentally (Qin et al. 2010) and can elicit greater magnitude and more specific responses (Sheth et al. 2016) than synthetically prepared treatments. In contrast to probiotics, microbiome engineering leverages complex microbial communities by engineering and transferring entire microbiomes, including unknown or unculturable bacterial strains, without prior knowledge of host-microbe interactions by selecting microbiomes based on host phenotype (Mueller & Sachs 2015).

As complex dynamic interactions among microbes in an *in-situ* microbial community are difficult to manipulate, only a few studies have so far tried to engineer native microbiome communities. Swenson et al. (2000) first engineered the *Arabidopsis thaliana* rhizosphere microbiome to increase and decrease shoot biomass by inoculating ten successive *Arabidopsis* selection rounds with the microbiome of plants with the highest or lowest above-ground biomass in the preceding round (Swenson et al. 2000). Panke-Buisse et al. (2015) expanded this application by selecting on late and early flowering-time under nutrient stress and demonstrating that the engineered microbiomes could influence flowering-time in additional *Arabidopsis* strains, as well as another related plant. Importantly, Panke-Buisse et al. (2015) evaluated microbiome composition through 16S amplicon sequencing, clearly illustrating that the microbiome evolved in response to host selection. However, Mueller and Sachs (2015) proposed the use of random-selection lines—where the propagated microbiome is randomly chosen from replicates—as the gold standard for experimental controls in microbiome engineering experiments, even while admitting that they greatly increase experimental effort. Previous microbiome engineering studies relied on sterile media transfers as negative controls, although Mueller et al. (2016) also incorporated a fallow-soil control for the presence of naturally occurring microbes. Notably, none of these studies used random-selection controls, which account for independent microbial evolution that may otherwise confound results. Furthermore, microbiome engineering experiments have, to the best of our knowledge, not yet been attempted with animal models.

In this study, we performed a microbiome engineering experiment incorporating random-selection lines in an animal model for the first time. We chose the fruit fly, *Drosophila melanogaster*, as a model for microbiome selection because of its relatively quick generation times (Trinder et al. 2017) and its simple core gut microbiome community (< 30 major species), which are largely commensals acquired from the environment (Erkosar et al. 2013; Blum et al. 2013). Furthermore, the functions and interactions of core members in the fly gut microbiome have been analyzed in detail (Broderick & Lemaitre 2012; Engel & Moran 2013); microbial interactions have been implicated in fly development, immunity, mating, response to external infection, and aging (Charroux & Royet 2012; Gould et al. 2018). Taking advantage of the fact that the microbiome affects fly development time (Shin et al. 2011; Ridley et al. 2012), we attempted to select for a microbiome that speeds up fly eclosion in sugar-starved flies and flies fed a high-sugar diet. Over the course of four selection rounds, we propagated the microbiome from vials with fast-eclosing flies and saw a significant decrease in fly eclosion times. However, there was no difference between selected treatments and random-selection controls. Our results emphasize the need for proper controls in microbiome evolution experiments and suggest that independent selection pressures on the microbiome may sometimes dominate in microbiome selection experiments.

## Materials and Methods

### Fly maintenance and phenotyping

The *Drosophila melanogaster* strain, Canton S, was used in this experiment because it has been kept inbred since its collection in the early 20th century (Stern 1943), which minimizes potential for host evolution over repeated experimental cycles (Emborski & Mikheyev 2019). Stock flies were reared on standard media with 45 g of cornmeal/100 g of sugar in an environmentally controlled incubator on a 12 hr:12 hr light/dark schedule. For the no-sugar diet, sugar and cornmeal were removed, whereas the high-sugar diet was prepared with 160 g of Glucose and no cornmeal. Sterile eggs were acquired by mating three-day old stock flies in egg collection cages with grape juice agar and yeast for 24 hours. The eggs were sterilized by gently rinsing 2× with a solution of double distilled water and 50% bleach for 30-120 seconds (Obadia et al. 2018; Newell & Douglas 2014).

As most eclosions take place during the day, the developmental time of the flies were assessed by recording the number of newly eclosed flies every hour during the 12 hr light period, from 9:00 AM to 9:00 PM. Eclosion times were recorded for three days, starting from the eclosion of first fly. To maintain fly genotype, flies were discarded after eclosion. Overall, 10,850 flies were phenotyped in the experiment.

### Indirect selection of the microbiome

We used the experimental protocol suggested by Mueller and Sachs (2015) for one-sided artificial selection on microbiomes (Figure 1). Twenty-four hour-old stock flies were transferred to fresh no-sugar and high-sugar diet vials (step 1 of Figure 1) for 3 days to make sure that all flies developed into sexually mature adults and that females had mated. The three day-old adult flies were transferred to fresh treatment media for 24 hours to lay eggs. These eggs, of approximately the same age, established the original microbiome community in high-sugar and no-sugar treatment diets.

**Figure 1.**
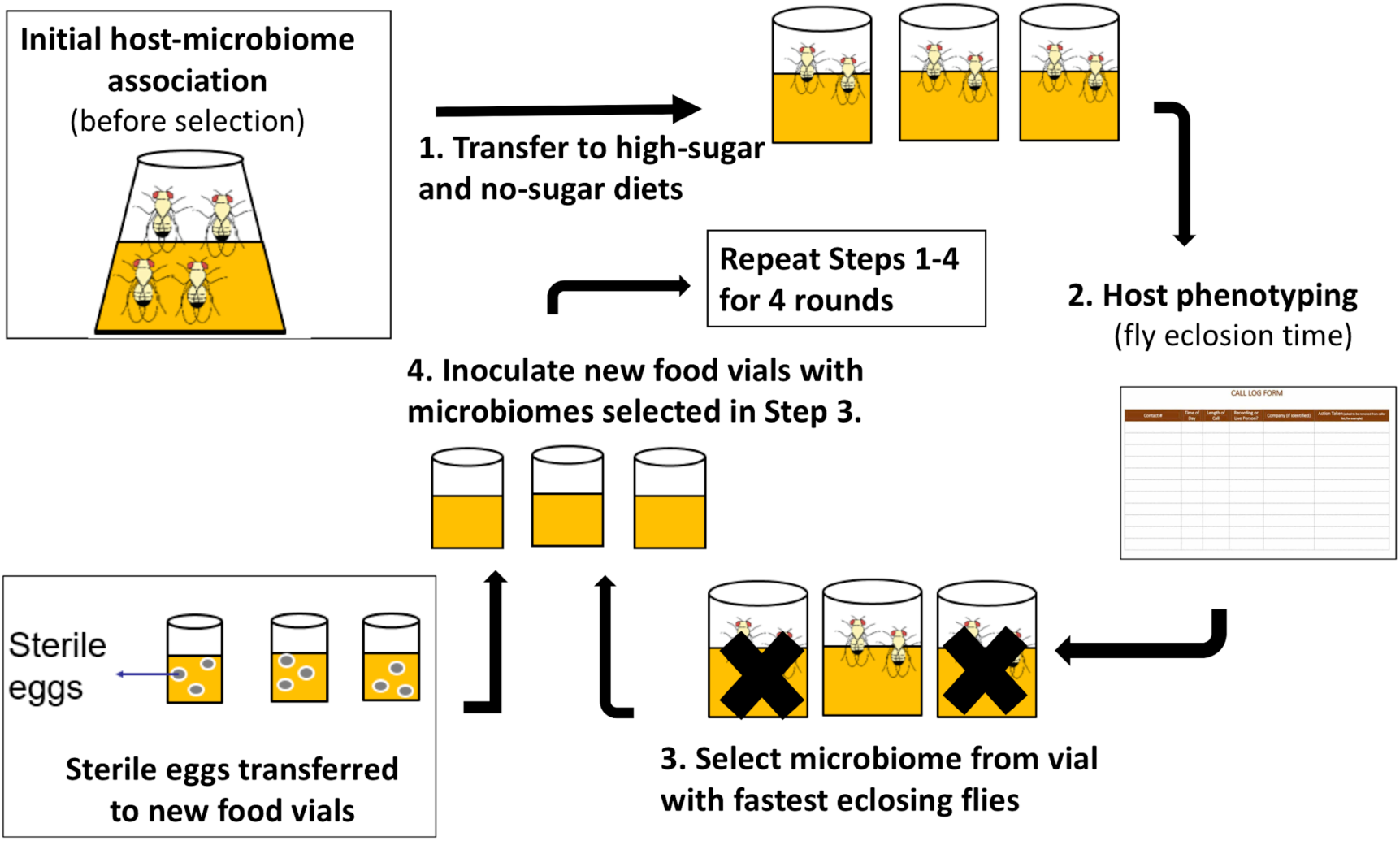
Schematic of experimental design for indirect selection of trait-associated microbiome in fruit flies. Following the experimental design suggested by Mueller and Sachs (2015), stock flies laid eggs in either high-sugar or no-sugar diets (step 1) and the microbiome from the fastest eclosing flies was propagated to the next selection round (steps 2-4). To keep host genotype constant, sterile eggs from stock flies were used in each selection round (step 4). There were ten parallel lines in each treatment, which were split into three sub-replicates at each round of selection. Random-selection lines were simultaneously maintained in high- and no-sugar media as experimental controls.

Each treatment (selection/high-sugar, selection/no-sugar, no-selection/high-sugar, no-selection/no-sugar) was initiated with 10 parallel lines, each of which was split into three sub-replicates at each selection round (30 vials per treatment). For selection treatments, the microbiome from the sub-replicate with the shortest mean eclosion time was selected to inoculate each of the three sub-replicates for that line in the next selection round (step 3 of Figure 1). For no-selection treatments, sub-replicates were randomly chosen for microbiome propagation to next round. Microbiome transfer was accomplished by passing the top layer (∼1 mm) of the fly food media through a 70 µm-mesh-size cell strainer (Fisher Scientific, cat no. 08-771-19) to remove any dead flies, unfertilized eggs or larvae and then equally distributing the strained media to the three sub-replicates of the corresponding line in next round. The top 1 mm of media was chosen as it is most likely to consist of native fly microbiome from the parent feces (Wong et al. 2015). Autoclaved spatulas were used for each food transfer to prevent any cross-contamination between lines. To ensure that the genetic pool of the flies remained constant and only the microbiome evolved, a spatula of sterilized eggs from the stock flies was aseptically transferred to each vial using an autoclaved spatula (step 4 of Figure 1). A spatula of media from vials chosen for propagation to the next round in selection and no-selection treatments was stored at −80°C for 16S rDNA sequencing. The selection procedure was repeated for a total of four rounds.

### 16S rDNA analysis

DNA extraction from media was performed using the DNeasy Blood and Tissue kit (QIAGEN, Hilden, Germany) following manufacturer’s protocols. Library preparation was done using the “16S Metagenomic Sequencing Library Preparation” protocol (Illumina) with 5% PhiX control added as an internal control for low diversity libraries. The libraries were sequenced by the Okinawa Institute of Science and Technology (OIST) sequencing section on the Illumina MiSeq platform with 2×250-bp v2 chemistry. The reverse read quality was too poor to join paired-end reads, however, and analysis was carried out on demultiplexed single-end sequences in QIIME 2 (v2017.11, Bolyen et al. 2018). The Divisive Amplicon Denoising Algorithm (DADA) was utilized through the DADA2 plug-in in QIIME 2 to quality-filter sequences, remove chimeras, and construct the Amplicon Sequence Variant (ASV) feature table(Benjamin J. Callahan et al. 2016). Taxonomic assignments were given to ASVs by importing SILVA 16S representative sequences and consensus taxonomy (release 128, Quast et al. 2013) to QIIME 2 and classifying representative ASVs using the naive Bayes classifier plug-in (Bokulich et al. 2018). The feature table, taxonomy, and phylogenetic tree were then exported from QIIME 2 to the R statistical environment (RC Team 2013) and combined into a Phyloseq object (McMurdie & Holmes 2013). To reduce the effects of uncertainty in ASV taxonomic classification, we conducted the analysis at the microbial ‘genus’ level. Prevalence filtering was applied to remove low-prevalence ASVs with less than 1% prevalence in order to decrease the possibility of data artifacts affecting the analysis (Callahan et al. 2016). Sequence counts were converted to relative abundance to normalize for varied library size and Weighted Unifrac (Lozupone et al. 2011) distances were computed between samples. Significance testing for distances between treatment groups was accomplished with the adonis function (Permutational Multivariate Analysis of Variance) in the Vegan R package (Oksanen et al. 2015), as well as the DESeq 2 pipeline implemented in phyloseq (Love et al. 2014).

### Statistical Analysis and Data Accessibility

Data were analyzed using the R statistical software (version 3.4.0; RC Team 2013). Exploratory analysis of the data to examine the fly eclosion time between no- and high sugar diets in each round were performed using tidyr (Wickham & Henry 2019) and ggplot2 packages (Wickham 2010). Linear mixed modelling was done to observe the effects of diet, round and selection on fly eclosion time using nlme package (Pinheiro et al. 2019) and visualized by Effects package (Fox et al. 2019).

All data and code necessary to reproduce the statistical tests, the main figures and tables are available on github (https://github.com/MikheyevLab/drosophila-microbiome-selection), including an interactive online document for the R-based analysis: https://mikheyevlab.github.io/drosophila-microbiome-selection/. Sequence data has been deposited into NCBI SRA database under the accession number PRJNA555001.

## Results

### Fly eclosion time is unaffected by artificial selection

To examine if diet, selection and round leads to faster fly eclosion time, we used linear mixed effects models, which allow for testing nested random effects and within-group variation. We used round, diet and artificial selection as fixed effects, and lines with sample replicates nested within them as random effects. We observed significant contribution of diet and round on fly eclosion time, but artificial selection did not affect the fly phenotype either as a main effect or an interaction (Figure 2, Table 1). Flies in high-sugar diets took longer to eclose than those in the no-sugar diet, but eclosion time decreased in both diets.

**Table 1.** Summary of mixed-effect model testing the effect of all the experimental parameters on fly eclosion time. While diet and selection round had strong effects on eclosion time, selection did not (also see Figure 2).

| *Predictors* | *Estimates* | *CI* | *p* |
| --- | --- | --- | --- |
| Intercept (High-sugar diet, No-selection control) | 292.96 | 288.36 – 297.57 | **<0.001** |
| Round of selection | -2.59 | -3.95 – -1.23 | **<0.001** |
| No-sugar diet | -50.00 | -56.29 – -43.70 | **<0.001** |
| Selection treatment | 3.58 | -3.21 – 10.37 | 0.302 |
| Round of selection × No-sugar diet | 0.81 | -1.06 – 2.68 | 0.397 |
| Round of selection × Selection treatment | -0.85 | -2.83 – 1.14 | 0.402 |
| No-sugar diet × Selection treatment | -1.52 | -10.57 – 7.53 | 0.742 |
| Round of selection × No-sugar diet × Selection treatment | 0.65 | -2.03 – 3.33 | 0.634 |
| N<sub>line</sub> | 10 |  |  |
| N<sub>replicate</sub> | 199 |  |  |
| Observations | 10850 |  |  |
| Marginal R<sup>2</sup> / Conditional R<sup>2</sup> | 0.779 / 0.834 |  |  |

**Figure 2.**
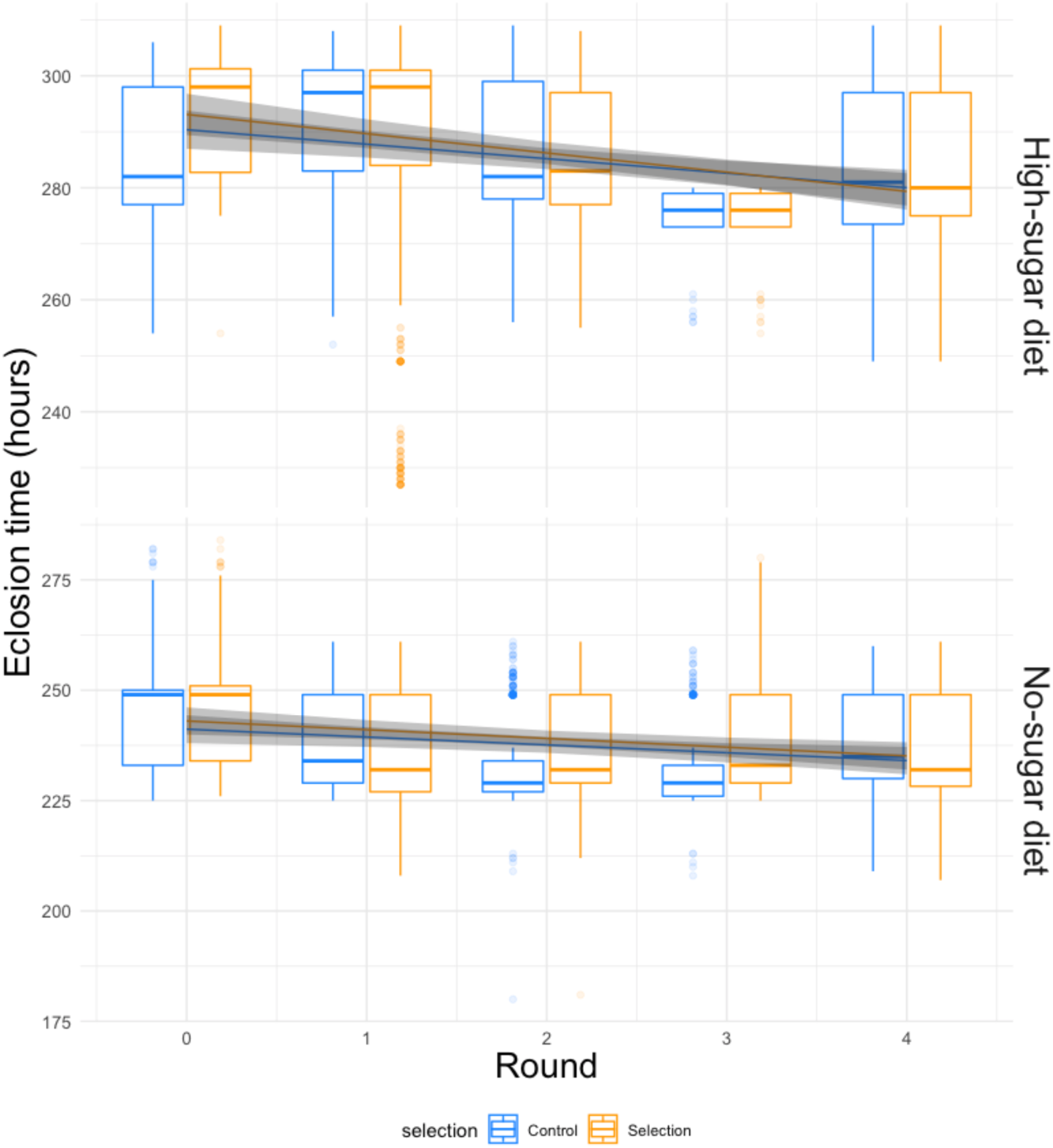
Phenotypic evolution over the course of the experiment. Box plots of raw data, and lines with 95% confidence intervals showing the fit of the mixed-effect linear model for the round by diet by selection interaction term (Table 1). In both diets, fly eclosion times decreased significantly over the course of the experiment. The difference in mean eclosion times between the first and last round of selection was 7.5±1.2 (S.E.) hours for the no-sugar diet and 12.1±1.2 (S.E.) hours for the high-sugar diet. However, selection had no effect and the rate of decrease was not different for control *vs*. selected flies. Rather that being driven by experimentally enforced selection, changes in phenotype were caused by independent evolution by the microbiome.

### 16S analysis of microbiome composition

We performed 16S amplicon sequencing of microbiome from both selected and no-selection control media that was chosen for propagation to the next round in each diet. We sequenced the V3/V4 hypervariable region of the 16S rRNA gene using the MiSeq v2 platform which generated an average of 175,522 reads per sample. These reads were analyzed using the DADA2 (Callahan et al. 2016) pipeline implemented in QIIME 2 (Bolyen et al. 2018). ASVs with low prevalence (< 0.01) were removed and alpha-diversity was measured by Shannon-diversity Index that accounts for both species abundance and evenness (Willis & Martin 2018). The association between bacterial alpha-diversity and artificial selection regime was tested via the adonis function in vegan R package (Oksanen et al. 2015), with alpha-diversity as dependent variable and diet, round, selection pressure as explanatory variables. The alpha-diversity varied with both diet and round. It increased in each successive round for both selected and non-selected vials, but it was more pronounced in no-sugar *vs*. high-sugar diet (Figure 3).

**Figure 3.**
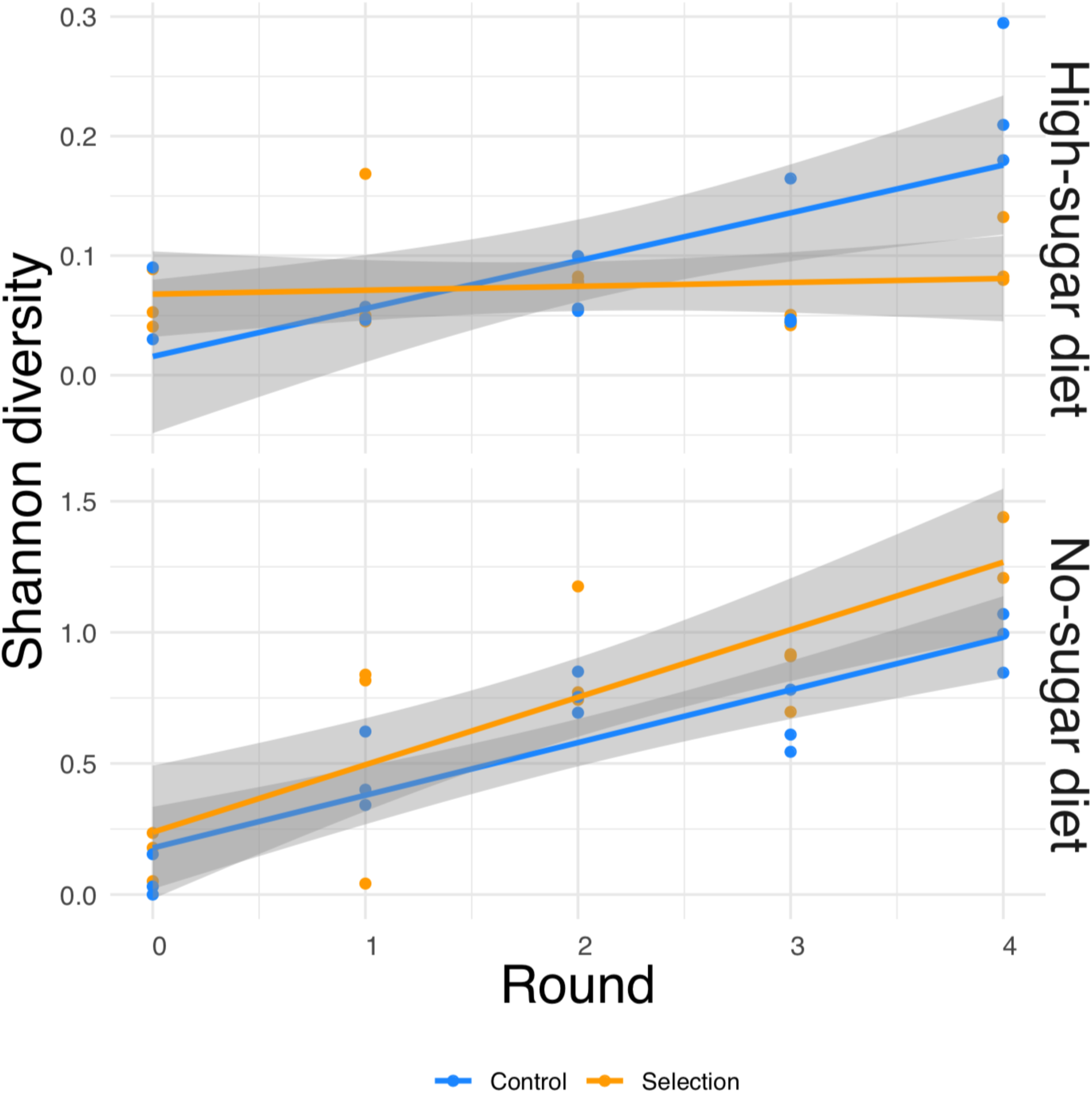
Alpha-diversity based on Shannon index for the media microbiome in high- and no-sugar diets over four rounds of artificial selection. Diversity was higher in the no-sugar diet (see also Figure 4), and increased over time (Table 2).

In general, the media contained low bacterial diversity, as reported previously (Blum et al. 2013). The microbial communities were dominated by Acetobacter initially, although other taxa increased in frequency, with a significant increase in alpha diversity over time in the no-sugar diet (Table 2, Figures 3 and 4). An adonis analysis of the UniFrac distances between microbial communities found a significant change over round for the no-sugar diet (F = 6.15, p = 0.0004), but not for the high-sugar diet (F = 0.62, p = 0.73), consistent with alpha diversity and compositional differences (Figures 3 and 4). We could not detect specific genera that systematically changed over the course of the experiment in either media using linear models implemented in DESeq2.

**Table 2.**
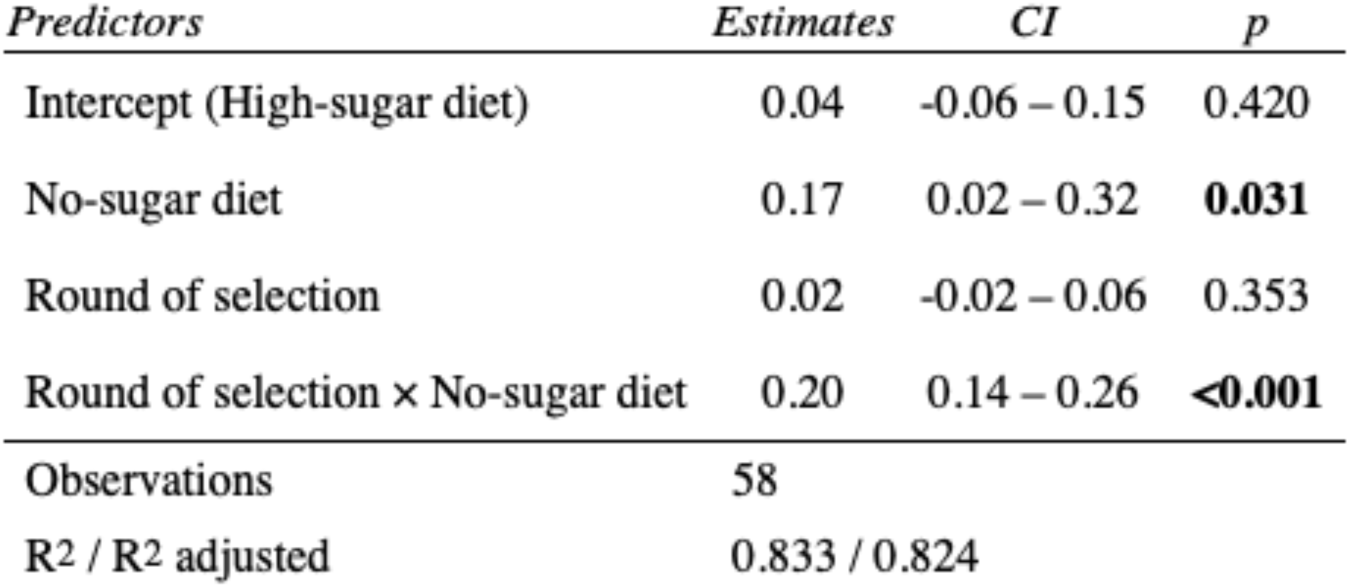
Shannon diversity index as a function of diet and selection round. Because there was no effect of selection (Table 1, also see adonis analysis), selection and no-selection treatments were combined to increase power. The alpha diversity was lower in the high-sugar diet (Figures 3 and 4), and increased over time in the no-sugar diet (Figure 3).

| *Predictors* | *Estimates* | *CI* | *p* |
| --- | --- | --- | --- |
| Intercept (High-sugar diet) | 0.04 | -0.06 – 0.15 | 0.420 |
| No-sugar diet | 0.17 | 0.02 – 0.32 | **0.031** |
| Round of selection | 0.02 | -0.02 – 0.06 | 0.353 |
| Round of selection × No-sugar diet | 0.20 | 0.14 – 0.26 | **<0.001** |
| Observations | 58 |  |  |
| R<sup>2</sup> / R<sup>2</sup> adjusted | 0.833 / 0.824 |  |  |

**Figure 4.**
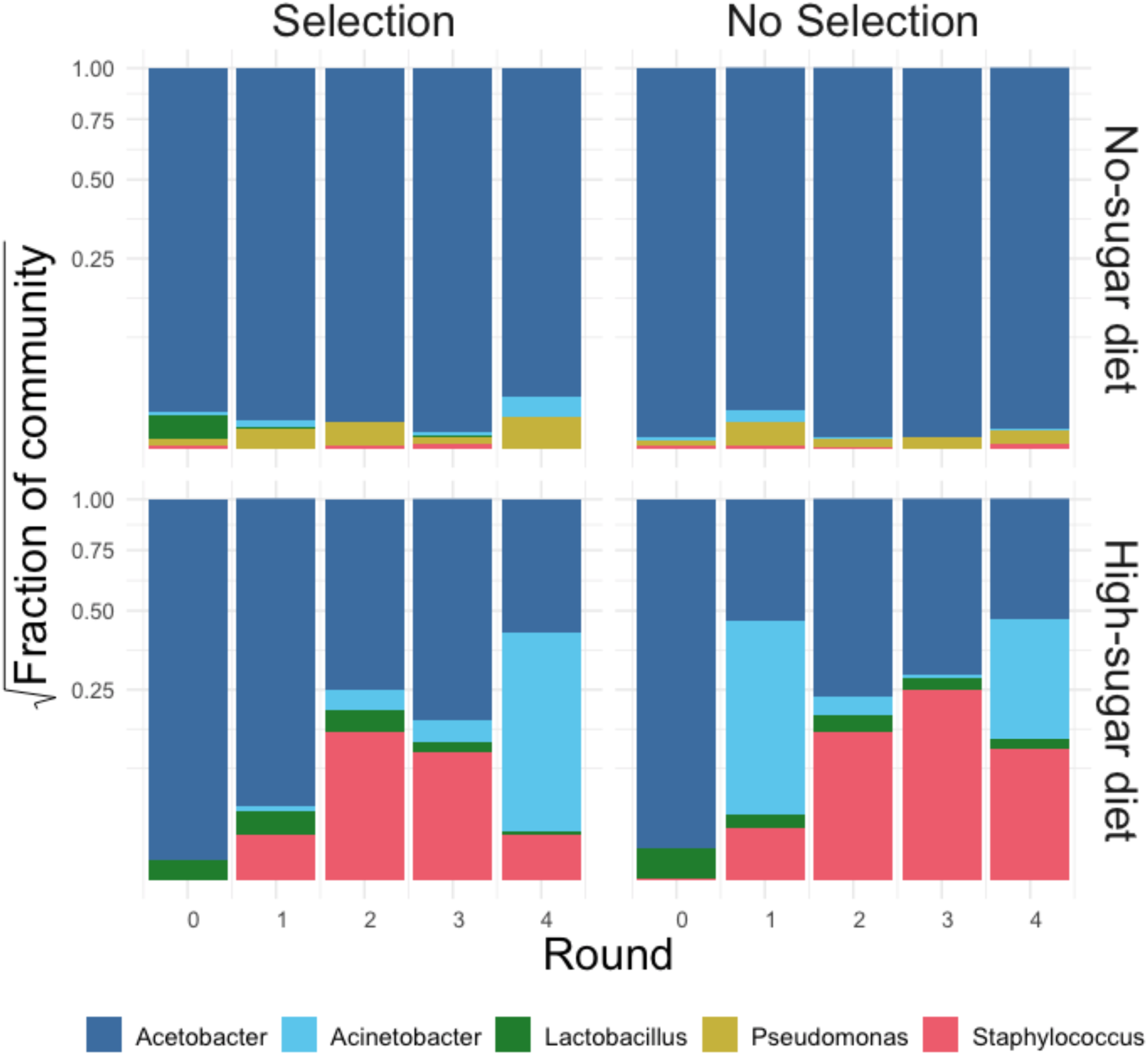
Stacked bar plot of bacterial genus-level relative abundance in the media over the course of the experiment. The compositional changes in community structure over time were only significantly different for the no-sugar diet, compared to the high-sugar diet (see also alpha diversity plots in Figure 3). The data are aggregated across three replicates in each condition. This suggests that the microbiome of the two media evolved differently, despite producing similar phenotypic results (Figure 2).

## Discussion

We attempted to apply microbiome engineering to increase fruit fly development rate by propagating the microbial community associated with fast development over four rounds of selection (Figure 1). We observed substantial increases in developmental rates over the course of the four rounds of selection in two different dietary media with no- or high-sugar content. However, selection for more rapid eclosion had no effect on either the developmental rate or its change over the course of the experiment (Figure 2). 16S amplicon sequencing showed that the microbial composition did indeed change over time, with a general increase in alpha diversity, particularly in the no-sugar diet, but it too was unaffected by selection (Figures 3 and 4). Thus, independent microbial evolution in the media swamped any signal of experimentally induced selection. However, by chance, the host phenotype changed in the same direction as our selection pressure. Only by using a random-selection line could we detect that the entirety of the observed effect was incidental.

To the best of our knowledge, this is the first study to examine host mediated indirect selection of microbiome in an animal model. The fruit fly is an excellent model for microbiome manipulations. It is an open symbiotic system, meaning that the internal and external microbes are similar (Wong et al. 2015; Blum et al. 2013). As a result, the processes affecting the microbiome can be complex, including a mixture of ecological, evolutionary and social interactions (Kaltenpoth et al. 2014; Mueller et al. 2005). For instance, behavior of individual flies, such as regurgitation and fecal deposition in the food, tunneling, and allo-coprophagy (consumption of conspecifics’ feces) lead to an exchange of symbionts among group members reared in the same media (Wong et al. 2015; Körner et al. 2016; Goodrich et al. 2014; Chandler et al. 2011). This aspect of fly biology motivated the media transfer in our experiment. However, since the inoculating flies came from a common stock, the indial microbiome diversity may have been low, with less variation available for subsequent artificial selection. Yet, the microbiome did evolve over subsequent rounds, with significant phenotypic effects on the flies.

It is well-known that the microbiome affects fly nutrition and development, particularly by interacting with the amount of fat (triglycerides) in the host; axenic individuals have a longer developmental period (Ridley et al. 2012; Newell & Douglas 2014; Ma et al. 2015; Shin et al. 2011; Storelli et al. 2011). However, the bacterial species have unique effects on the host. It is likely that our measures of bacterial relative abundance and community diversity metrics (Figures 3, 4) cannot fully capture the complexity of bacterial interactions with the host. This is exemplified by the fact that we could not detect any significant changes in the bacterial community associated with high-sugar media, though the composition of no-sugar media’s did change significantly according to our measurements. Yet, both media had comparable decreases in fly eclosion times. It is, therefore, possible that phenotypic changes resulted from effects of lower-frequency strains, or perhaps from other factors, such as chemical compounds produced by the microbes in response to each other. In complex systems, such as microbial communities, substantial phenotypic variation may be due to interactions between its components, which play a role in facilitating community-level selection (Williams & Lenton 2007).

### Efficacy of experimental selection *vs*. independent evolution by the microbiome

Microbial evolution experiments typically apply discrete rounds of selection (Swenson et al. 2000; Panke-Buisse et al. 2015; Mueller et al. 2016). For example, our experiment had four distinct rounds, where media from fast-developing vials was transferred to the next round. However, media microbiome evolved continuously in between rounds of selection and not necessarily in ways that we wanted or could effectively control. For example, to be passed to the next round of selection, microbes had to aggressively colonize fresh media and compete amongst themselves for resources, but not in a way that negatively affected fly larvae. Because there are many microbial generations within selection rounds, these parallel selection pressures may dominate the evolutionary response with significant effects on the host phenotype, as appears to have happened in our experiment.

This sort of independent evolution of the microbiome may significantly limit the utility of microbiome engineering. However, the topic has received relatively little theoretical or empirical attention (though see Williams & Lenton (2007)). One key implication is for the design of controls during microbiome evolution studies. Randomly selected control lines allow the microbiome to evolve in the same way, except for the experimentally enforced selection. However, these controls are extremely time- and labor-consuming. Alternative options, such as constant inoculation from a preserved microbiome source, or null inoculations, have been proposed as efficient alternatives (Mueller & Sachs 2015). Even the fallow-soil control used by Mueller et al 2016, which is a substantial methodological advance over typically used sterile controls, doesn’t take into account possible interactions between the microbiome and the plant and how they might evolve. Experimental designs with other control strategies do not provide the same level of control over microbial evolution as does the random-selection control. For example, using constant or null controls in our experiment would have lead us to erroneously infer that the microbiome evolved in response to experimental selection.

Along similar lines, we cannot exclude the possibility that the host has changed in the course of the experiment. Strictly controlling the host population (*e*.*g*., in a glycerol stock or seed bank) is not possible with fruit flies. In retrospect, it would have been desirable to confirm stability of eclosion times in the source population at the beginning and the end of the experiment. Changes in the host population appear a less likely explanation for the observed data, given the magnitude of change seen in the experiment – about half a day earlier eclosion in the course of four rounds (Figure 2). First, the fly stocks were inbred and genetically homogeneous, minimizing the possibility of evolutionary changes (Emborski & Mikheyev 2019). Second, they were kept in a stable environment with controlled temperature, humidity, photoperiod and diet. Third, eggs were surface sterilized to prevent the introduction of additional microbes to the experiment. Nonetheless, even because the most stable-seeming environments, such as glycerol stock or seed banks may experience change over time (*e*.*g*., due to freezer malfunctions or fungal rot), ideally both host and microbiome changes should be controlled in the course of microbiome engineering experiments. Therefore, we strongly recommend that studies introduce this ‘host-stability’ control.

In conclusion, the findings show that artificial selection is not significantly correlated with fly phenotype or microbiome. This was made possible due to the use of random-selection controls to measure selection pressure. The lack of significant correlation of selection might be driven by factors independent of host-mediated artificial selection. Any future prospects in artificial engineering of host microbiome to select desirable host phenotype would require selection regimes that are stronger than microbial evolution.

## Acknowledgements

We are grateful to Carmen Emborski and Ulrich Mueller for help in designing this study. We thank Takakazu Yokokura and Cecelia Lu for providing invaluable training and assistance in fly-rearing and egg collection. We thank the OIST DNA sequencing section – Onna, Okinawa, for carrying out the sequencing. This work has been funded by the Okinawa Institute of Science and Technology Graduate University.

